## Supplementary material for "Rapid assembly of SARS-CoV-2 genomes reveals attenuation of the Omicron BA.1 variant through NSP6"

**A**

| | Overhang | | | | |
|---|---|---|---|---|---|
| | 5′ | 3′ | nt | nt | ORF |
| **F1** | ATTA | GTGC | 1 | 2721 | ORF1a (nsp1&2) |
| **F2** | GTGC | GAGA | 2718 | 5454 | ORF1a (nsp3) |
| **F3** | GAGA | GTAA | 5451 | 8556 | ORF1a (nsp3) |
| **F4** | GTAA | TCTA | 8553 | 11846 | ORF1a (nsp4–6) |
| **F5** | TCTA | TGCA | 11843 | 15090 | ORF1a (nsp7-11), ORF1ab (nsp12) |
| **F6** | TGCA | GCTG | 15087 | 18043 | ORF1ab (nsp12&13) |
| **F7** | GCTG | CAAT | 18040 | 21564 | ORF1ab (nsp14–16) |
| **F8** | CAAT | GAAC | 21561 | 25390 | S |
| **F9** | GAAC | ACGA | 25387 | 27891 | ORF3a/b, E, M, ORF6, ORF7a/b |
| **F10** | ACGA | AAAA | 27888 | 29908 | ORF8, N, ORF9b/c, ORF10 |

**B**

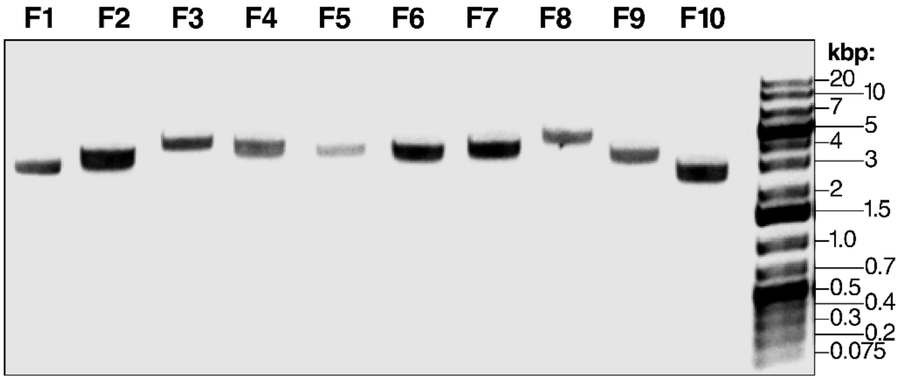


**Supplementary Figure 1. SARS-CoV-2 fragment design and quality.**

(A) Coordinates of the 10 SARS-CoV-2 fragments and the sequences of the 5′ and 3′ overhangs when the fragments were digested by BsaI. Coordinates are based on the WA1 sequence.

(B) Agarose gel electrophoresis of the 10 fragments from A after PCR amplification and clean up. 200 ng of each fragment was loaded on the gel.

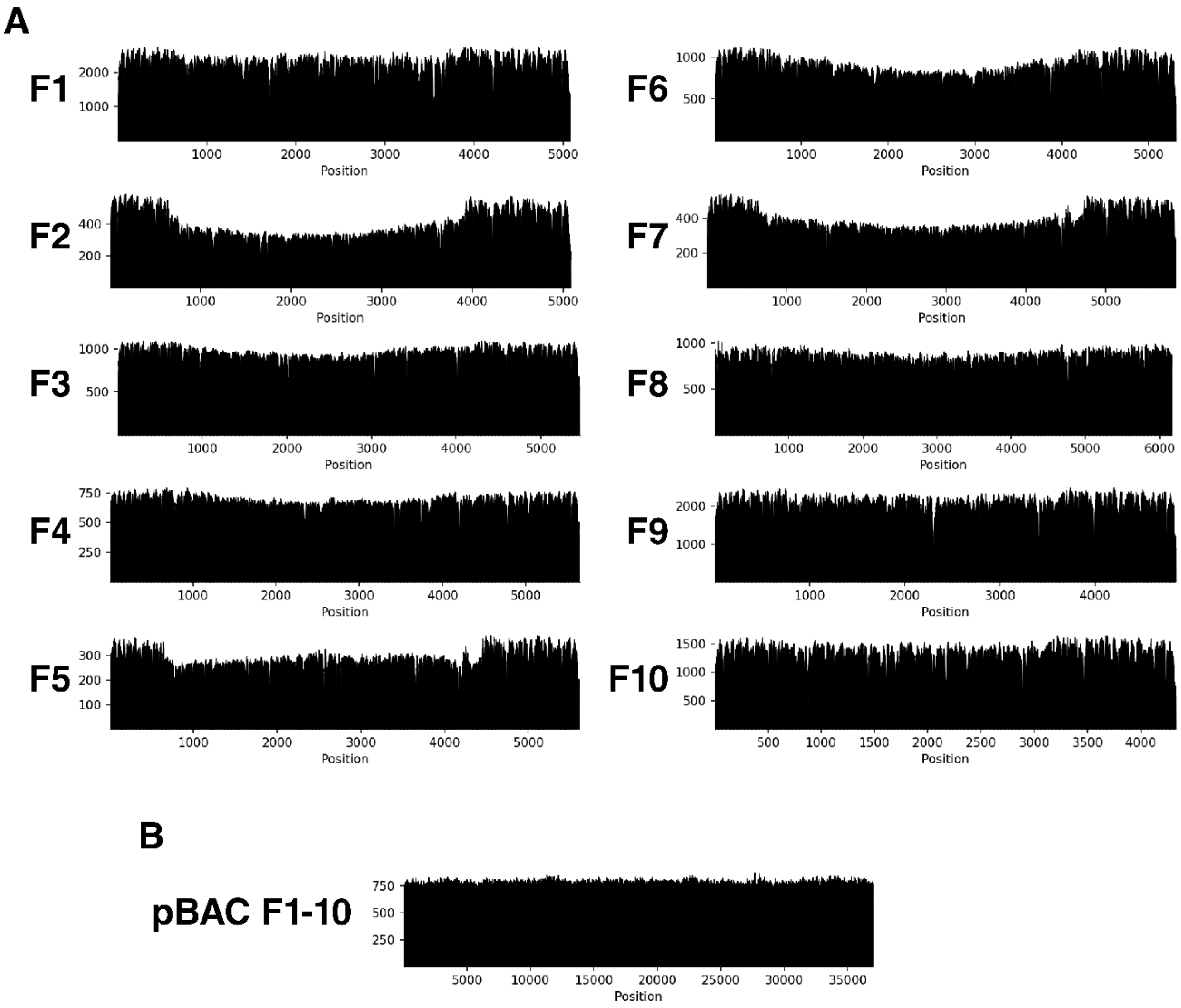


**Supplementary Figure 2. Nanopore sequencing of the 10 SARS-CoV-2 genome fragment plasmids and the pBAC SARS-CoV-2 plasmid.**

(A) Representative coverage (# reads mapped per base) plots for each of the 10 SARS-CoV-2 genome fragment plasmids. Sequencing was done through Primordium Labs whole plasmid sequencing service.

(B) Representative coverage (# reads mapped per base) plot of the pBAC SARS-CoV-2 plasmid. Sequencing was done through Primordium Labs whole “Large” plasmid sequencing service.

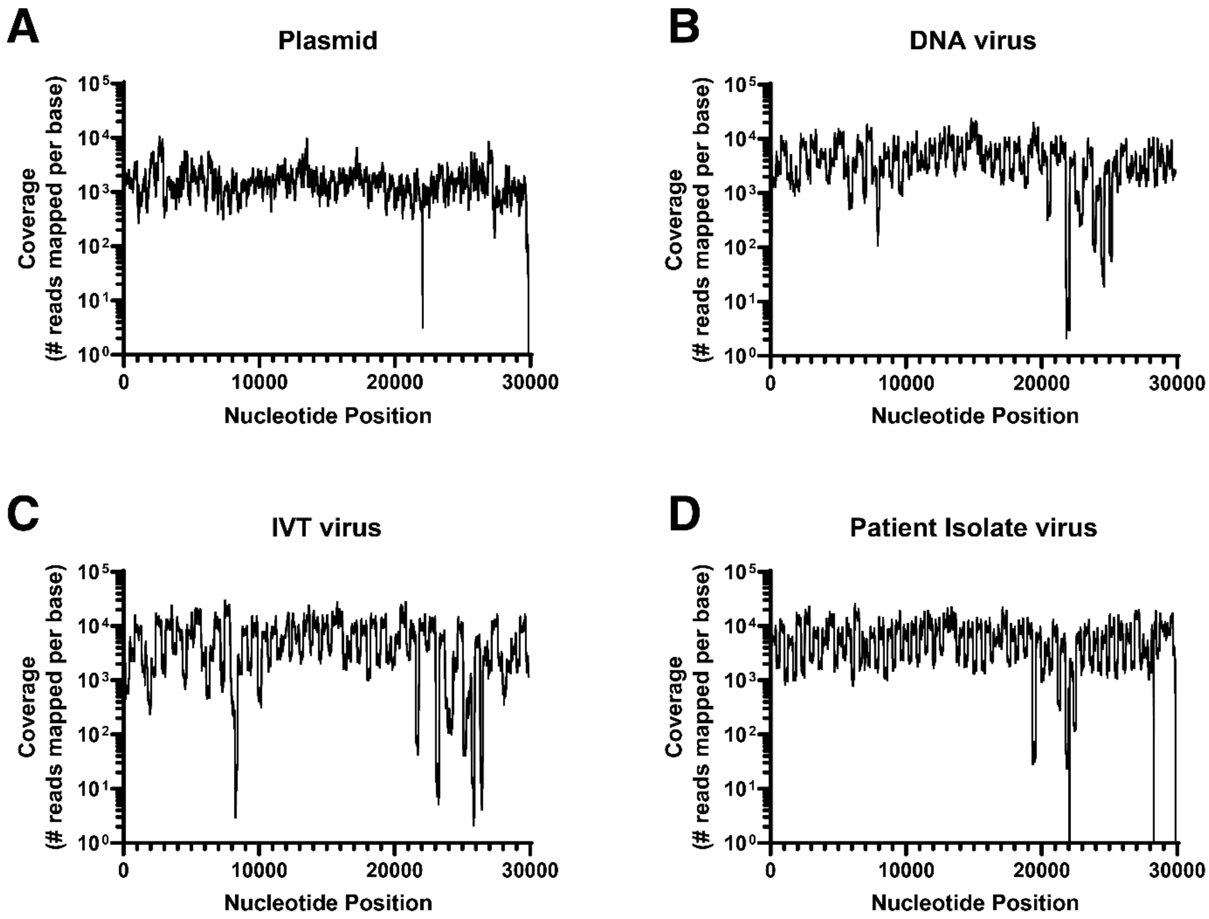


**Supplementary Figure 3. NGS sequence verification of pBAC SARS-CoV-2 plasmid and patient isolate, DNA- and RNA-launched viruses.**

Representative coverage (# reads mapped per base) plots of the plasmid used to generate the viruses in Fig. 2 (A), including DNA- (B) and RNA-launched (C) viruses and the patient isolate virus (D). All sequencing was done using the ARTIC Network's protocol.

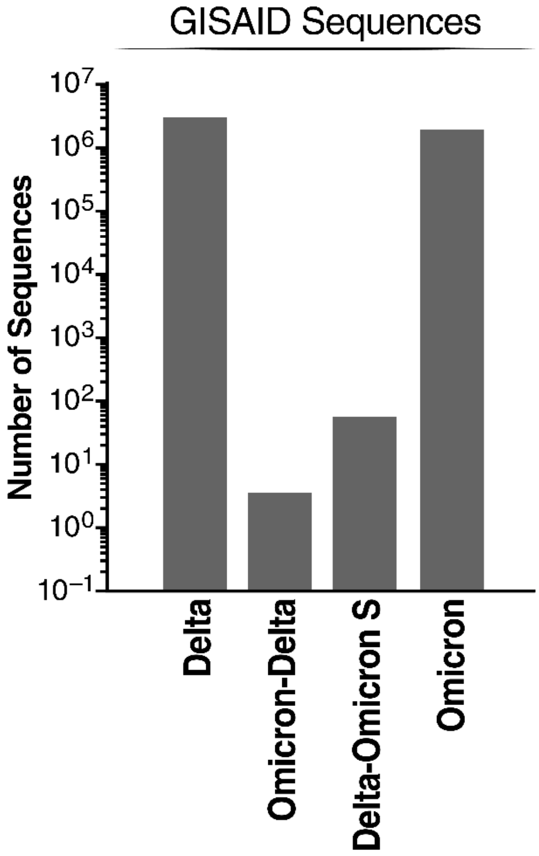


**Supplementary Figure 4. Abundance of SARS-CoV2 Delta, Omicron, and Delta-Omicron recombinant sequences.**

Sequences were extracted from the GISAID database using key mutations specific to each of the Delta and Omicron variants (see Fig. 3 for more details). Omicron-Delta indicates an ORF1ab and ORF2-10 recombinant.

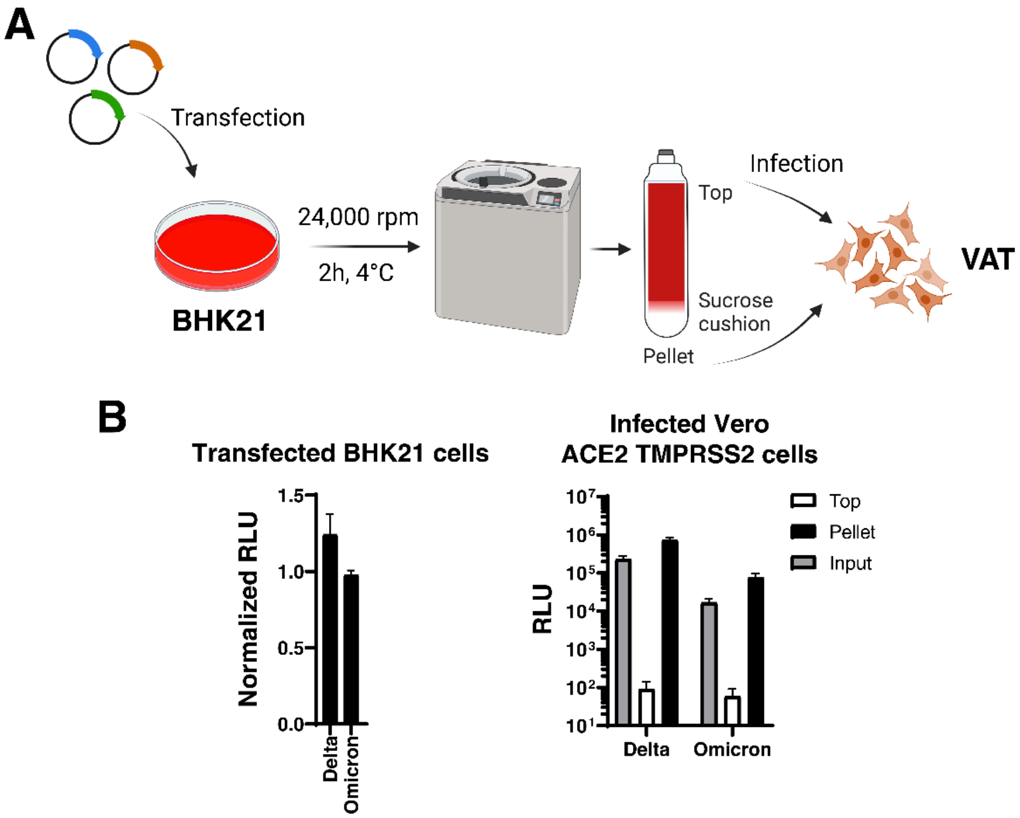


**Supplementary Figure 5. Concentration and validation of SARS-CoV-2 replicon-containing particles.**

(A) Workflow for the concentration and validation of SARS-CoV-2 replicon-containing particles. VAT: Vero cells stably expression ACE2 and TMPRSS2.

(B) Luciferase readout from transfected BHK21 and infected Vero ACE2 TMPRSS2 cells of Delta and Omicron replicons. Data shown are average ± SD of three independent transfections and concentration experiments.

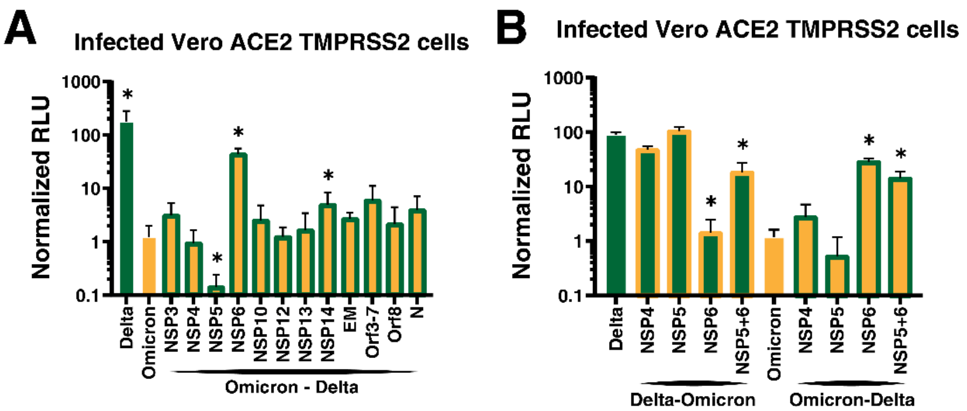


**Supplementary Figure 6. Omicron mutations in NSP6 reduce viral RNA replication.**

(A) Luciferase readout from infected Vero ACE2 TMPRSS2 cells with supernatant from BHK21 cells transfected with Delta, Omicron, and Omicron-Delta recombinants replicons as indicated and an Omicron spike and nucleocapsid expression vectors. Average of three technical replicates ± SD are shown, and pairwise comparisons were made relative to the Omicron variant by two-sided Student's T-test.

(B) Luciferase readout from infected Vero ACE2 TMPRSS2 cells with supernatant from BHK21 cells transfected with Delta, Delta-Omicron recombinants, Omicron, and Omicron-Delta recombinants replicons as indicated and an Omicron spike and nucleocapsid expression vectors. Average of three technical replicates ± SD are shown, and pairwise comparisons were made relative to the Omicron variant by two-sided Student's T-test.
